# Adolescent THC impacts on mPFC dopamine-mediated cognitive processes in male and female rats

**DOI:** 10.1101/2024.04.12.588937

**Authors:** Maricela X. Martinez, Vanessa Alizo Vera, Christina M. Ruiz, Stan B. Floresco, Stephen V. Mahler

## Abstract

**Rationale:** Adolescent cannabis use is linked to later-life changes in cognition, learning, and memory. Rodent experimental studies suggest Δ^9^-tetrahydrocannabinol (THC) influences development of circuits underlying these processes, especially in the prefrontal cortex, which matures during adolescence.

**Objective:** We determined how 14 daily THC injections (5mg/kg) during adolescence persistently impacts medial prefrontal cortex (mPFC) dopamine-dependent cognition.

**Methods:** In adult Long Evans rats treated as adolescents with THC (AdoTHC), we quantify performance on two mPFC dopamine-dependent reward-based tasks—strategy set shifting and probabilistic discounting. We also determined how acute dopamine augmentation with amphetamine (0, 0.25, 0.5 mg/kg), or specific chemogenetic stimulation of ventral tegmental area (VTA) dopamine neurons and their projections to mPFC impacts probabilistic discounting.

**Results:** AdoTHC sex-dependently impacts acquisition of cue-guided instrumental reward seeking, but has minimal effects on set-shifting or probabilistic discounting in either sex. When we challenged dopamine circuits acutely with amphetamine during probabilistic discounting, we found reduced discounting of improbable reward options, with AdoTHC rats being more sensitive to these effects than controls. In contrast, neither acute chemogenetic stimulation of VTA dopamine neurons nor pathway-specific chemogenetic stimulation of their projection to mPFC impacted probabilistic discounting in control rats, although stimulation of this cortical dopamine projection slightly disrupted choices in AdoTHC rats.

**Conclusions:** These studies confirm a marked specificity in the cognitive processes impacted by AdoTHC exposure. They also suggest that some persistent AdoTHC effects may alter amphetamine-induced cognitive changes in a manner independent of VTA dopamine neurons or their projections to mPFC.

## Introduction

Cannabis is one of the most widely used drugs among adolescents, and its availability is increasing around the world. Human studies show that early exposure to cannabis, and especially its main psychoactive constituent Δ^9^-tetrahydrocannabinol (THC), is associated with later-life cognitive impairments, and increased risk for psychiatric disorders including schizophrenia and addiction (Curran et al. 2016; Ehrenreich et al. 1999; Jenni et al. 2017; Malone et al. 2010; Murray et al. 2022; Rubino and Parolaro 2016; Schneider 2008; Volkow et al. 2016). However, in humans it is difficult to dissect whether THC exposure causes these associations, or whether early cannabis use and long-term deficits both result instead from other underlying comorbidities or risk factors. Rodent models are thus essential for establishing casual effects of THC on the developing adolescent brain.

Neurodevelopmental disruptions persisting long after adolescent cannabis use are plausible because adolescence is a dynamic critical period for structural and functional brain remodeling, especially in late-developing structures like the prefrontal cortex (PFC) (Andersen 2003; Casey et al. 2000). Some of this age-dependent plasticity seems to involve the endocannabinoid system, with dynamic changes in cannabinoid receptors (CBRs) and endocannabinoids (ECB) occurring across adolescence (Bara et al. 2021; Ellgren et al. 2008; Heng et al. 2011; Lee et al. 2016; Simone et al. 2022). Might THC, which also acts via CBRs, disrupt this age-dependent ECB signaling system and thus leave long-lasting consequences on the brain? If so, the adolescent-developing PFC (Peters et al. 2022; Scheyer et al. 2023; Spear 2000), and its dopaminergic inputs from ventral tegmental area (VTA), which are actively innervating during this period (Hoops and Flores 2017; Manitt et al. 2011; Reynolds et al. 2018), are a likely candidate for cognition-relevant neurodevelopmental insults caused by adolescent THC (Molla and Tseng 2020; Renard et al. 2017a; Renard et al. 2017b).

The adult PFC is crucial for purposeful, goal-directed behaviors driven by the ability to flexibly converge our internal states, like reward motivation, with outside external information about contexts, cues, and rules (Miller 2000; Miller and Cohen 2001; Ott and Nieder 2019). Executive functions like working memory, attention, rule shifting, and decision-making require PFC-dependent cognitive control. Rodent studies show adolescent cannabinoid drug exposure can cause persistent deficits in working memory (De Melo et al. 2005; O’Shea et al. 2004; Quinn et al. 2008; Schneider and Koch 2003), social cognition (O’Shea et al. 2004; Renard et al. 2017a; Renard et al. 2017b; Zamberletti et al. 2014), and cognitive flexibility (Egerton et al. 2005; Gomes et al. 2015; Jacobs-Brichford et al. 2019; Szkudlarek et al. 2019) that may depend upon PFC.

Furthermore, dopamine in PFC plays a major role in decision making, working memory, cognitive flexibility, and goal-directed behaviors (Floresco 2013; Goldman-Rakic 1995; Goto et al. 2007; Seamans and Yang 2004). PFC dopamine dysfunction has also been implicated in schizophrenia symptoms (Davis et al. 1991; Howes and Kapur 2009), and there is a clear link between recent cannabis use and onset of psychosis in humans (Andreasson et al. 1987; D’Souza et al. 2004; Hambrecht and Hafner 2000), though it is not clear that this link is causal in nature (Fergusson et al. 2005; Henquet et al. 2005; Sewell et al. 2009). Since VTA dopamine neuron axons actively infiltrate mPFC during adolescence (Hoops and Flores 2017; Manitt et al. 2011; Reynolds et al. 2018) and are thus subject to disruption by THC exposure, and since THC impacts dopamine signaling at several key nodes in reward and salience circuits (Behan et al. 2012; Corongiu et al. 2020; Ferland et al. 2023; Renard et al. 2017a; Renard et al. 2017b), it is plausible that adolescent THC might exert some of its neurodevelopmental disruptions by impacting the function of cognitive flexibility-relevant dopaminergic inputs to mPFC.

We therefore explored this possibility in a series of experiments quantifying effects of a well-characterized, translationally relevant adolescent THC exposure protocol in rats (Ruiz et al. 2021a; Ruiz et al. 2021b; Torrens et al. 2022; Torrens et al. 2020) upon adulthood mPFC dopamine-dependent cognition.

## Materials and Methods

### Subjects

Long Evans rats (*n* = 29 males, 25 females) were used for set shifting, probabilistic discounting and amphetamine experiments, and transgenic TH:Cre (*n* = 19 males, 9 females) and wildtype littermates *(n=* 14 males, 4 females) were used for chemogenetic experiments. Rats were pair or triple-housed in same-sex groups at weaning (postnatal day; PD21), in a temperature, humidity, and pathogen-controlled colony under a 12:12hr reverse light/dark cycle. Water was provided *ad libitum* and food was restricted to ∼85% of free-feeding weight during behavioral testing, starting at ∼PD70. Experiments were approved by University of California Irvine’s Institutional Animal Care and Use Committee and were conducted in accordance with the NIH Guide for the Care and Use of Laboratory Animals.

### Drugs

THC was provided in ethanol by the NIDA Drug Supply Program. For injection, THC was prepared fresh each day; ethanol is evaporated under N_2_, and THC is reconstituted in 5% Tween-80 in saline, with heat and sonication, to 2 ml/kg for intraperitoneal (IP) injection. D-amphetamine hemisulfate salt (amphetamine) was attained from Sigma and mixed in saline at 0.5 mg/ml for injection. Clozapine-N-oxide (CNO) was obtained from the NIDA Drug Supply Program, stored at 4°C in powder aliquots with desiccant, protected from light. For systemic injection, CNO (5 mg/kg) was dissolved daily for IP injection in 5% dimethyl sulfoxide (DMSO; Sigma Aldrich) in 0.9% saline. For microinjections, CNO (1mM; 0.5 µl/side over 60s) was dissolved in artificial cerebrospinal fluid (Fisher) with 0.5% DMSO, stored in aliquots at −20°C, and thawed/vortexed just before use.

### Adolescent THC and Washout

All rats received daily IP injections of THC (5 mg/kg) or vehicle (VEH; 5% Tween-80 in saline) from PD30-43, followed by a washout period of 21+ days, allowing full THC clearance in both sexes (Lee et al. 2022) prior to behavioral testing in adulthood (**Fig. 1a**).

**Fig. 1.**
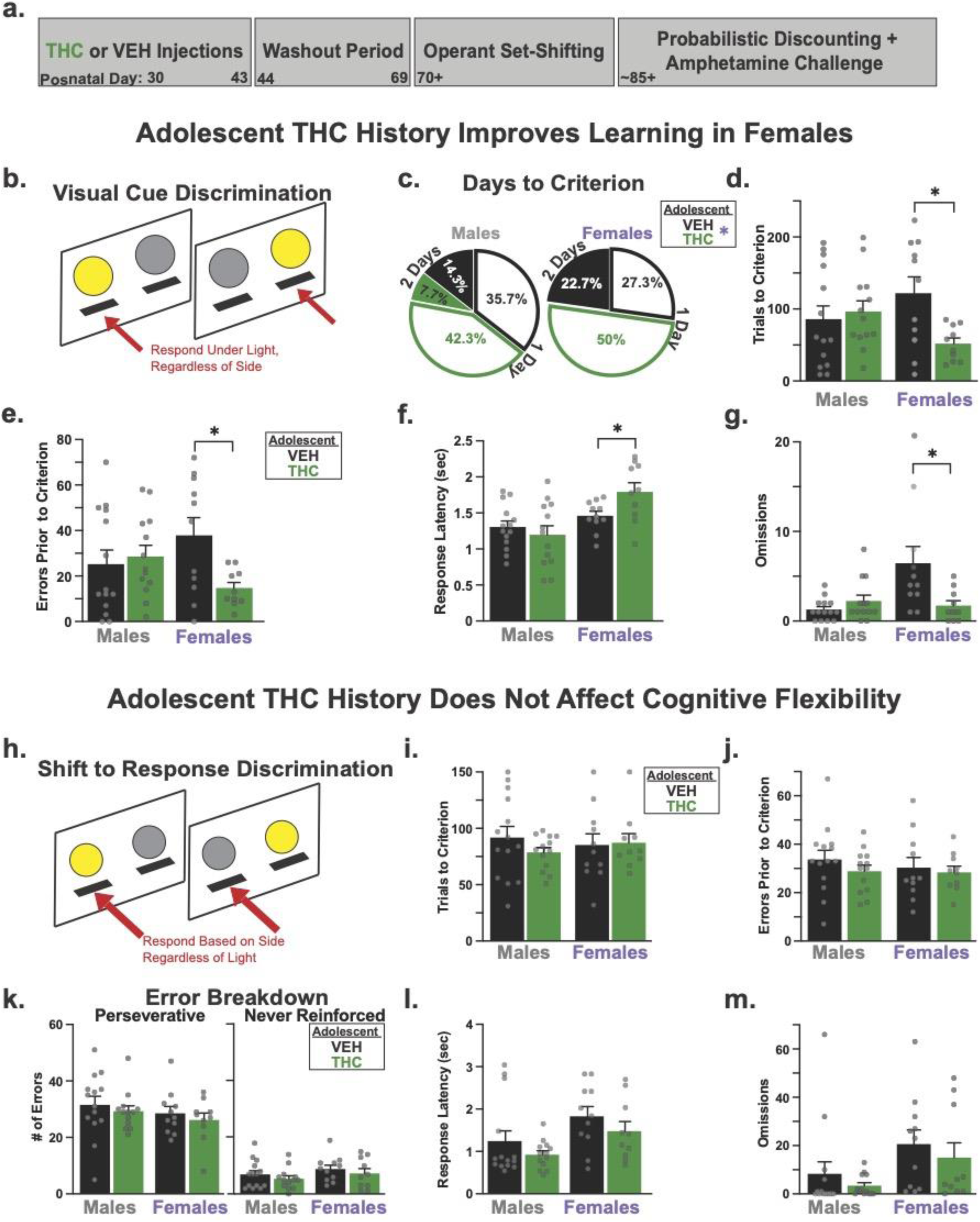
Adolescent THC history selectively impacts adulthood learning and cognition. **a)** Experimental timeline. **b)** Schematic of the visual cue discrimination task. **c)** AdoTHC rats (green wedges) were more likely than AdoVEH rats (black wedges) to acquire the visual cue discrimination task to criterion in only one training session (unfilled wedges), rather than requiring 2 sessions to acquire (filled wedges). Both sexes showed similar patterns. **d, e)** AdoTHC females took fewer trials to meet criterion, and made fewer errors when learning visual cue discrimination than AdoVeh females, no such effects were seen in males. **f, g)** AdoTHC females took longer to respond, and made fewer errors than AdoVEH females during cue discrimination training, without effects in males. **h)** Schematic of the subsequently tested strategy set-shifting task. **i)** AdoTHC did not alter the number of trials to learn the new rule to criterion in either sex. Likewise, AdoTHC did not alter in either sex **j)** the number of errors, **k)** the types of errors, **l)** latency to respond, or **m)** omitted trials during set shifting training. Individual rats shown as grey dots in each graph: AdoVEH (*n* = 14 males, 11 females), AdoTHC (*n* = 13 males, 10 females). X^2^ *p*\* < 0.05 and repeated measure two-way ANOVA, Sidak post hoc: *p*\* < 0.05. Data presented as mean + SEM.

### Experimental Design

Following adolescent THC/VEH treatment, 48 adult (PD 70+) rats (*n* = 29 males, 21 females) underwent strategy set-shifting training, followed by training on probabilistic discounting until a group displayed stable levels of choice for 3 consecutive days, determined with previously established criteria (St. Onge and Floresco 2009). Thereafter they underwent 3 counterbalanced amphetamine challenge tests, each (0, 0.25, 0.5 mg/kg IP) delivered 5 min before behavioral testing. Following the challenge rats were retrained for at least 2 days before receiving their next challenge.

For chemogenetic experiments, another adolescent THC/VEH-treated group (*n* = 33 males, 13 females; *n* = 28 TH:Cre+,18 wildtype (WT)) underwent stereotaxic VTA virus injection of a Cre-dependent hM3Dq vector at ∼PD65, and intra-mPFC bilateral cannulae implantation at least 45 days later, at least 8 days prior to the first microinjection. Following recovery from surgery, they were trained on the probabilistic discounting task to stability, then subjected to a series of counterbalanced tests, again with re-stabilization training occurring between them. First, rats received 4 counterbalanced IP injections of the DREADD agonist CNO (5 mg/kg) or VEH, 30 min prior to behavioral testing. Both CNO and VEH were administered twice on separate days, and data was combined between both tests for analysis. Next, they underwent 4 additional counterbalanced tests of probabilistic discounting, each held 5 min after intra-mPFC microinjections of CNO or VEH. Results from intra-mPFC VEH and CNO tests were again averaged to increase reliability of findings. For choice data, the raw number of risky choices in each block made on both VEH or CNO tests were summed and divided by the total number of choices made in those respective blocks on the two VEH/CNO tests, thereby factoring out trial omissions. Win-stay/lose shift values were combined in a similar manner, dividing the sum total of “stays” or “shifts” by the sum total of “wins” and “losses” on the two respective tests. Latency and omission values were averaged across the tests.

### Behavioral Methods

#### Operant Boxes

Training and testing took place in Med-Associates rat operant conditioning chambers (30.5 x 24 x 21 cm; St Albans, VT) within sound-attenuating boxes, equipped with two retractable levers with white lights above them, and white house light.

#### Operant Pretraining

Methods for both strategy set shifting and reward probabilistic discounting tasks closely followed prior reports (Floresco et al. 2008; St. Onge and Floresco 2009). Rats were first homecage-habituated to highly palatable, 45 mg banana-flavored reward pellets (Bio-Serv catalogue #: F0059), then given 2 days of magazine training in the operant boxes, where they received 38 pellets at variable intervals over a 60 min session. They were next sequentially trained to press each of two levers to receive a pellet over 2-5 days. On each lever training day, a lever extended into the chamber (side counterbalanced across rats), and one pellet was delivered on a fixed ratio 1 (FR1) schedule. Once they reliably pressed 50+ times in 30 min on a lever, they were transitioned to learning to press the other lever on the subsequent day, and training continued until meeting this criterion. Thereafter, they entered the next phase of the task, in which each lever was periodically extended into the chamber throughout a 30 min session. Levers extended in a pseudorandom order such that there were 45 left-lever trials and 45 right-lever trials, but no more than two consecutive trials on which the same lever was extended. Each lever extension was accompanied by illumination of a house light signaling the start of a trial (90 trials/session), and trials occurred every 20s. If the rat pressed the lever within 10s of the start of a trial, the lever was retracted and a pellet was delivered, and the house light remained on for 4s. If the rat failed to press in 10s, the lever retracted and the house light extinguished until the next trial, and the trial was considered an omission. Importantly, cue lights present above each lever were never illuminated during this training phase. Rats were trained in this manner for at least 5 days, or until they omitted less than 5 trials per session. Since individual rats frequently display idiosyncratic lever position biases that can influence interpretation of subsequent behavior (Brady and Floresco 2015), and to minimize such impacts of stochastic side-preference on subsequent tests, lever side preference was next assessed using a published protocol (Brady and Floresco 2015; Floresco et al. 2008).

#### Visual Cue Discrimination Training

Next, rats were trained to respond on only one of the two levers extended on each trial—whichever was signaled as the correct response by illumination of a cue light just above it (**Fig. 1b**). Sessions began with both levers retracted and the house light off. Every 20s, one of the two stimulus cue lights was illuminated in a randomized order, 3s later both levers extended, and the house light turned on. If the rat pressed the lever that had a cue light illuminated above it within 10s of lever insertion, it received a pellet, the levers retracted, the stimulus light was extinguished and the house light remained illuminated for 4s. If the rat did not choose a lever in 10s, the trial ended and levers retracted until the next trial, and the trial was scored as an omission. This training proceeded for at least 30 trials, ending when rats either made 10 consecutive correct responses, or after 150 trials had elapsed. If rats did not achieve 10 consecutive correct responses, they received a second identical training session on the following day.

#### Strategy Set Shifting Test

After learning to follow a cue light to respond for reward in the prior training phase, rats then underwent a single 40 min session on which the response rule was suddenly shifted; a procedure analogous to the Wisconsin Card Sorting Task used in humans to aid diagnosis of PFC dysfunction (Owen et al. 1991; Pantelis et al. 1999). On this day, the non-preferred lever (determined in side-bias training described above) became the correct response, and pressing it during trials yielded a pellet and commencement of the next inter-trial interval (**Fig. 1h**). Light cues above one of the levers were presented just before and during trials as in the prior visual cue discrimination training, but now their location was irrelevant to the receipt of reward. Instead, rats needed to recognize that this old rule (follow the cue light) no longer worked, and that pressing of the previously less-preferred lever, regardless of cue light position, was now the correct strategy for obtaining reward. Trials continued until rats performed 10 consecutive correct responses, or after 150 trials had occurred. Errors during set-shift were categorized into two subtypes as in (Floresco et al. 2008): perseverative errors, where rats responded on the incorrect lever when the previously-relevant visual cue was illuminated above it, and never-reinforced (or non-perseverative) errors, where rats pressed the incorrect lever despite when the visual cue was illuminated above the correct lever.

#### Probabilistic Discounting

In this task rats chose between a lever that always delivers a small (1 pellet) “certain” reward, or another that delivers a large (4 pellet) “risky” reward, delivered at various probabilities throughout the 52.5 min session (**Fig. 2a**). The probability of receiving 4 pellets upon pressing the risky lever decreases in each of five sequential choice blocks during each session (100%, 50%, 25%, 12.5%, 6.25%). Each block began with 8 forced trials (35s apart), in which the houselight was illuminated and 3s later, one of the two levers extended one at a time for 10s (4 trials for each lever, randomized in pairs). This permitted rats to sample the probability of reward receipt upon pressing of the risky lever in the upcoming certain/risky lever choice phase of the block. During the subsequent choice phase of each block, both levers were extended, and after rats pressed one of them, both levers retracted. Certain lever presses always yielded 1 tasty banana pellet, and risky lever choices delivered 4 or 0 banana pellets, at a probability that varied across blocks. Failure to respond on a lever within 10s on any trial resulted in lever(s) retraction and extinguishing of the house light, and the trial was scored as an omission. Animals were trained on this task for 21 days, at which point choice behavior had stabilized across groups. For rats tested with systemic amphetamine during probabilistic discounting, training on this task commenced following strategy set shifting training and testing described above. Rats in chemogenetic experiments did not undergo visual cue discrimination or set shifting tests. Instead, these rats received pellet habituation, magazine training, initial FR1 training, and retractable lever training prior to training on the probabilistic discounting task.

**Fig. 2.**
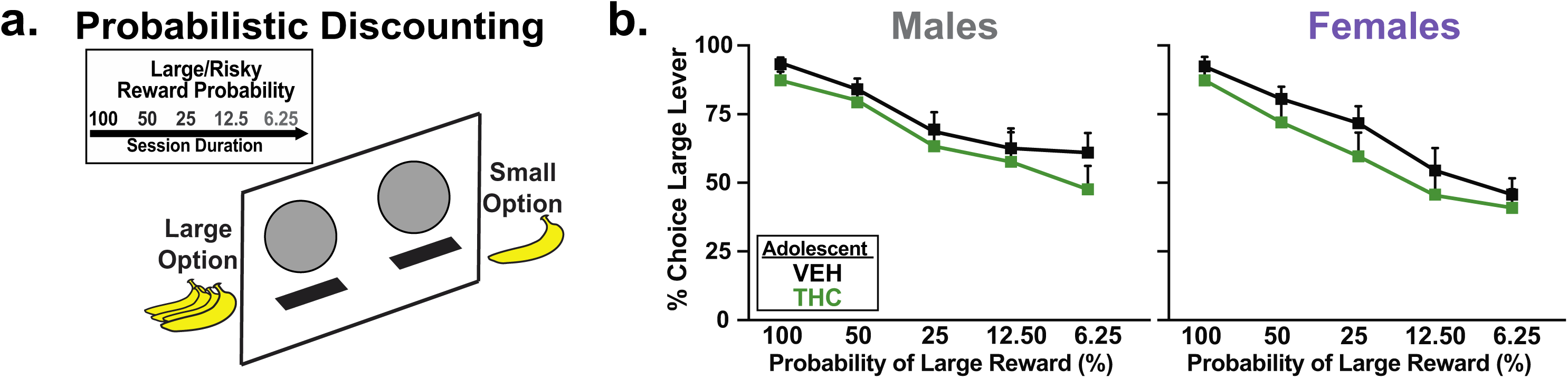
AdoTHC history does not affect baseline probabilistic discounting. **a)** Schematic of the probabilistic discounting task. Sessions consisted of 5 training blocks, in which 2 levers are presented, and rats must choose either a small reward lever that always delivers one pellet, or a large reward-delivering lever which becomes increasingly unlikely to deliver any reward as the session progresses. **b)** Data is shown for males and females, indicating that AdoTHC rats did not differ from their AdoVEH counterparts in probabilistic discounting, with both shifting from nearly exclusively preferring the large reward lever, but appropriately shifting away from it as it became less likely to deliver reward. AdoVEH (*n* = 14 males, 11 females), AdoTHC (*n* = 15 males, 8 females). Data represented as average within each probability block from three consecutive days of stable performance. Mean ± SEM.

For rats that received acute amphetamine challenges (0, 0.25, 0.5 mg/kg IP) on risky decision making, 3 counterbalanced tests were conducted on separate days, with re-training between tests to reestablish stable baseline responding. When rats in chemogenetic experiments achieved stable baseline responding, they similarly received 4 counterbalanced tests, 2 with CNO and 2 with VEH (data from both tests combined for analysis as described above), all injected IP 30 min prior to probabilistic discounting testing, with retraining between tests. After completing these systemic CNO/VEH tests, rats were implanted with mPFC cannulae, allowed to recover, and re-stabilized on discounting performance for 2+ days. They were then tested on the discounting task in 4 additional counterbalanced tests with intra-mPFC microinjections of CNO (1mM/0.5 µl; 2 tests) and VEH (2 tests).

### Chemogenetic Methods

#### Virus Surgery

Rats were anesthetized with ketamine (56.5 mg/kg) and xylazine (8.7 mg/kg) and given meloxicam (1.0 mg/kg) for pain prophylaxis. An AAV2 vector containing a Cre-dependent, mCherry-tagged hM3Dq excitatory DREADD (hSyn-DIO-hM3Dq-mCherry; titer: 6 x 10^12^ vg/ml; Addgene catalog #: 44361) was injected bilaterally into VTA (relative to bregma (mm): AP: −5.5, ML: ±0.8, DV: −8.1; 0.75 µL/hemisphere) using a Picospritzer and glass micropipette (Martinez et al. 2023). Injections occurred steadily over 1 min, and the pipette was left in place for 5 min after injection to limit spread. Both TH:Cre and WT rats were injected with the active hM3Dq DREADD virus. Colocalization of mCherry expression to TH+ VTA neurons was verified in each TH:Cre rat, and lack of mCherry expression was confirmed in each WT rat. At least 3 weeks elapsed between virus injections and the first CNO administration. In a second surgery held following behavioral training and systemic CNO tests (to maximize accuracy of placement) guide cannulae (22 ga, 2MM, Plastics One) were implanted bilaterally in the mPFC (relative to bregma (mm): AP: 2.7, ML: ±1, DV: −3.1) to allow intra-mPFC CNO injection upon VTA dopamine neuron axon terminals, and occluded with steel stylets between tests.

#### Histological Validation

Following behavioral testing, chemogenetic rats were perfused with chilled 0.9% saline and 4% paraformaldehyde, brains were cryoprotected in 20% sucrose-azide, and they were sectioned coronally at 40 μm. VTA virus expression was amplified with mCherry immunohistochemistry, and expression verified to be in dopamine neurons via co-staining of tyrosine hydroxylase. VTA sections were then blocked in 3% normal donkey serum PBST, tissue was incubated overnight at room temperature in rabbit anti-DSred (Clontech; 1:2500) and mouse anti-TH (Immunostar; 1:2000). After washing, sections were incubated in dark at room temperature for 4 hours in AlexaFluor-donkey anti-Rabbit 594 and donkey anti-Mouse 488 (Thermo Fisher Scientific; 1:500). Sections were mounted and cover slipped (Fluoromount; Thermo FisherScientific), mCherry/TH expression was imaged at 10X on a Leica DM4000 epifluorescent scope, and expression or lack thereof in VTA was verified in each animal. To verify cannula placement within mPFC, sections were nissl stained with cresyl violet, and cannula tracks were mapped using a rat brain atlas (Paxinos and Watson 2006).

### Data Analysis

Effects of adolescent treatment (AdoTX: THC or VEH) and Sex (M or F) on learning and cognition employed 2 x 2 ANOVA and Sidak posthoc tests. Effects of amphetamine (0 x 0.25 x 0.5mg/kg) on probabilistic discounting were tested with repeated measures ANOVA, and interactions of these acute treatements with AdoTX and Sex were examined by adding these between subjects variables to multivariate General Linear Model ANOVAs. Simple-main effects analyses conducted after observing staticially significant interactions included one-way ANOVA models using the appropriate error term from the overal multifactor analyses. Effects of CNO versus VEH were treated as within-subjects variable, while Genotype (TH:Cre x WT), AdoTX, and sex were treated as between subjects variables in multivariate General Linear Model ANOVAs. Prior to testing, rats were verified to display stable patterns of risky choice by examining discounting performance over at least two consecutive prior training days. Rats of both sexes were tested in chemogenetic experiments, but sample sizes for each sex were insufficient to allow formal analysis of this variable (Fig. S1). Six rats (*n* = 2 males, 4 females; *n* = 1 VEH, 5 THC) were excluded from set shifting analyses for failure to meet training critera for instrumental or visual cue rule performance, and nine rats (*n* = 9 females; *n* = 2 VEH, 7 THC) tested with amphetamine were excluded for failing to achieve stable performance on the probabilistic discounting task. Nine rats (*n* = 3 males, 6 females; *n* = 4 VEH, 5 THC) were excluded from the DREADD experiments for failure to stabilize on the probabilistic discounting, death, or inability to confirm virus expression.

## Results

### Effects of Adolescent THC on Strategy Set Shifting in Each Sex

#### Initial Instrumental Training

AdoTHC did not affect acquisition of instrumental food seeking behavior during initial training (no main effect of AdoTX: Fs_(1, 45)_ < 0.40, *ps* > 0.53), and acquisition was similar in both sexes (No main effect of sex: Fs_(1, 45)_ < 3.60, *ps* > 0.06; no Sex x AdoTX interaction: (Fs_(1, 45)_ < 1.12, *ps* > 0.30).

#### Visual Cue Discrimination

Rats were next trained to press whichever lever that had a cue light illuminated above it. AdoTHC exposed rats were more likely to learn the visual cue rule in one session rather than two, relative to AdoVEH rats (X^2^: *p* = 0.04). This was driven by AdoTHC females, as evidenced by these animals requiring fewer trials and making fewer errors to reach criterion performance of 10 consecutive correct choices (**Fig. 1c-g**, AdoTX x Sex interaction; trials to criterion: F_(1,44)_ = 5.31, *p* = 0.03; errors to criterion: F_(1,44)_ = 5.03, *p* = 0.03; Šídák’s posthoc for trials *p* = 0.02, errors *p* = 0.02). In addition, relative to controls, AdoTHC females were slower to respond (F_(1, 44)_ = 4.55, *p* = 0.04) but also made fewer omissions: (F_(1, 44)_ = 8.46, *p* = 0.01). In contrast, no differences were observed on these measures between AdoTHC males and controls (trials to criterion: *p* = 0.87; errors: *p =* 0.89; latency: *p* = 0.68, omissions: *p* = 0.72). More generally, we also saw some sex-differences on performance measures irrespective of treatment, as females were slower to respond (Sex: F_(1,44)_ = 13.02, *p* < 0.001) and omitted more trials (F_(1,44)_ = 5.60, *p* = 0.02) compared to males.

#### Strategy Set Shifting

Following training on the initial visual cue discrimination, rats were then trained to use an egocentric spatial rule (always press the left/right lever regardless of cue location), and tested in a single-session (Fig. 1i-m; Brady and Floresco 2015). AdoTHC had few effects on the ability to learn this new response rule with no change in either sex seen on the number of trials to reach criterion on the new rule (No main effect of AdoTX: F_(1,44)_ = 0.42, *p* = 0.52; or Sex: F_(1,44)_ = 0.01, *p* = 0.92; or interaction: F_(1,44)_ = 0.80, *p* = 0.38). Total errors to criterion were also unaffected (AdoTX: F_(1,44)_ = 0.97, *p* = 0.33; Sex: F_(1,44)_ = 0.32, *p* = 0.57; AdoTX x Sex interaction: F_(1,44)_ = 0.19, *p* = 0.66), as were errors of either a perseverative or never-reinforced subtype (AdoTX: F_(1,44)_ = 1.56, *p* = 0.22; Sex: F_(1,44)_ = 0.13, *p* = 0.72; AdoTX x Sex: F_(1,44)_ = 0.58 x 10^3^, *p* = 0.98). Additionally, we saw no effect of AdoTX on response latency or on omissions (AdoTX: Fs _(1,44)_ < 2.62, *ps* > 0.11; AdoTX x Sex: Fs _(1,44)_ < 0.01, *ps* > 0.94). Again, we saw that females took longer to respond (Sex: F_(1,44)_ = 7.52, *p* = 0.01) and omitted more trials (F_(1,44)_ = 6.20, *p* = 0.02) compared to males. Since visual discrimination learning was more efficient in AdoTHC females than in AdoVEH females, but shifting to a spatial rule was equivalent in both groups, we wondered if performance on rule #1 could have led to more robust or persistent learning that could have indirectly impacted performance on the set shift. We therefore performed a secondary analysis where we matched the performance of the two female AdoTX groups on the visual discrimination rule #1. We removed the 4 AdoTHC female rats that learned rule #1 quickest, and the 5 AdoVEH females that learned rule #1 slowest, thus yielding equivalent rule #1 performance in the retained rats from both groups (n=6/group; Errors: AdoVEH: M (SEM) = 17.83 (6.18) AdoTHC: M (SEM) = 19.00 (2.94); t(10) = 0.17, *p* = 0.87; Trials: AdoVEH: M (SEM) = 16.83 (22.83); AdoTHC: M (SEM) = 64.67 (9.58); t(10) = 0.17, *p* = 0.870). Additionally, in this subset, performance on the shift to rule #2 remained equivalent (Errors: AdoVEH: M (SEM) = 79.33 (16.07); AdoTHC: M (SEM) = 91.50 (13.04); t(10) = 0.22, *p* = 0.83; Trials: AdoVEH: M (SEM) = 27.17 (6.58); AdoTHC: M (SEM) = 28.83 (3.95); t(10) = 0.59, *p* = 0.57), rather than showing an AdoTHC-induced deficit as would be expected if rule #1 performance impacted subsequent shift to rule #2.

### Effects of Adolescent THC on Probabilistic Discounting in Each Sex

#### Acquisition of Probabilistic Discounting

The same rats were next trained on a probabilistic discounting task for 21 days. All rats had acquired stable performance by the last three days of training (no block X day interaction: F_(5.498,241.89)_ = 1.70, *p* = 0.13). No effect of AdoTX (F_(1, 44)_ = 0.82, *p* = 0.37), Sex (F_(1,44)_ = 0.12, *p* = 0.74), or AdoTX x Sex interactions (F_(1, 44)_ = 0.59, *p* = 0.45) were found, suggesting that rats of both sexes and AdoTX histories acquired the discounting task at similar rates. *Stable Probabilistic Discounting:* Following training, all rats exhibited stable and comparable performance on the task by the last three days, with decreasing choice of the high-reward lever as delivery of this reward became increasingly unlikely or “risky” (main effect of Block: F_(4,176)_ = 75.01, *p* < 0.001). No effect of AdoTX (F_(1,44)_ = 0.85, *p* = 0.36), Sex (F_(1,44)_ = 0.10, *p* = 0.75), or AdoTX x Sex interactions (F_(1, 44)_ = 0.62, *p* = 0.44) were found, suggesting that rats of both sexes and AdoTX histories displayed comparable levels of risky choice across blocks (**Fig. 2b**). Likewise, neither AdoTX nor Sex affected win stay or lose shift choice strategies (no main effect of AdoTX: Fs_(1,44)_ < 1.64, *ps* > 0.21; no main effect of Sex Fs_(1,44)_ < 0.55, *ps* > 0.46; no AdoTX x Sex Fs_(1,44)_ < 1.51, *ps* > 0.23; data not shown).

With respect to other performance measures, AdoTX did not affect choice latencies (no AdoTx: F_(1, 44)_ = 0.16, *p* = 0.69; no AdoTX X Sex interaction: F_(1, 44)_ = 0.1.29, *p* = 0.26). Additionally, we found that AdoVEH rats omitted more on the “riskier” probability blocks compared to AdoTHC animals (AdoTX x Sex x Block: F_(4, 176)_ = 3.95, *p* = 0.004, AdoTX x Block: F_(4, 176)_ = 4.39, *p* = 0.002; no AdoTx: F_(1, 44)_ = 0.94, *p* = 0.34), an effect that was particularly apparent in females (AdoTX x Block: F_(4, 68)_ = 3.19, *p* = 0.02; data not shown). Overall, we saw that females took longer to respond than males (Sex: F_(1,44)_ = 40.03, *p* < 0.0001) and omitted more trials (F_(1,44)_ = 63.83, *p* < 0.0001), especially in low-probability training blocks at the end of the session (Sex x Block: Latency: F_(4,176)_ = 4.57, *p* = 0.002; Omissions: F_(4,176)_ = 26.57, *p* < 0.0001).

#### Effects of Acute Amphetamine on Probabilistic Discounting

As previously reported (St. Onge et al. 2010), amphetamine increased choice of the large/risky option in a dose dependent manner when analyzed across both groups and sexes (**Fig. 3a**, amphetamine x Block interaction: F_(8,328)_ = 10.02, *p* < 0.0001; main effect of amphetamine: F_(2,82)_ = 9.88, *p* < 0.0001). However, the effects of different doses of amphetamine varied as a function of AdoTX treatment and sex. Analysis of the choice data also revealed significant amphetamine x AdoTX (F_(2,82)_ = 3.33, *p* = 0.04) and amphetamine x AdoTX x Sex interactions (F_(2,82)_ = 3.20, *p* = 0.05). Partitioning this latter interaction by AdoTX group revealed that for control rats, amphetamine exerted different effects on choice in males vs females (amphetamine x Sex: F_(2,44)_ = 3.73, *p* = 0.03). Post-hoc comparisons showed that, in males, amphetamine increased risky choice following treatment with the 0.5 mg/kg dose (*p* < 0.02) but not the lower, 0.25 mg/kg dose (*p* = 0.19). In contrast, the 0.5 mg/kg dose had more deleterious effects on choice in females, reducing risky choice in the high probability blocks and increasing it in the lower ones, while the 0.25 mg/kg dose induced a modest increase in risky choice in the latter block. This yielded an overall lack of effect of amphetamine in control females (**Fig. 3b**, amphetamine: F_(2,18)_ = 0.57, *p* = 0.57). On the other hand, amphetamine induced a more reliable and pronounced increases in risky choice in both males and females AdoTHC animals, with analysis of these data producing significant main effects of amphetamine treatment (**Fig. 3a,b**, F_(2,38)_ = 12.07, *p* < 0.0001) in the absence of an interaction with the sex factor (F_(2,38)_ = 1.56, *p =* 0.22). When collapsed across sex, post hoc comparisons showed that both the 0.25 and 0.5 mg/kg dose increase risky choice (both *p*s < 0.05), although inspection of Fig. 3b indicates that the effect of the 0.25 mg/kg dose was driven primarily by females. From these data, we conclude that AdoTHC treatment makes rats more sensitive to the ability of amphetamine to increase risky choice, and this effect appears to be more prominent in females.

**Fig. 3.**
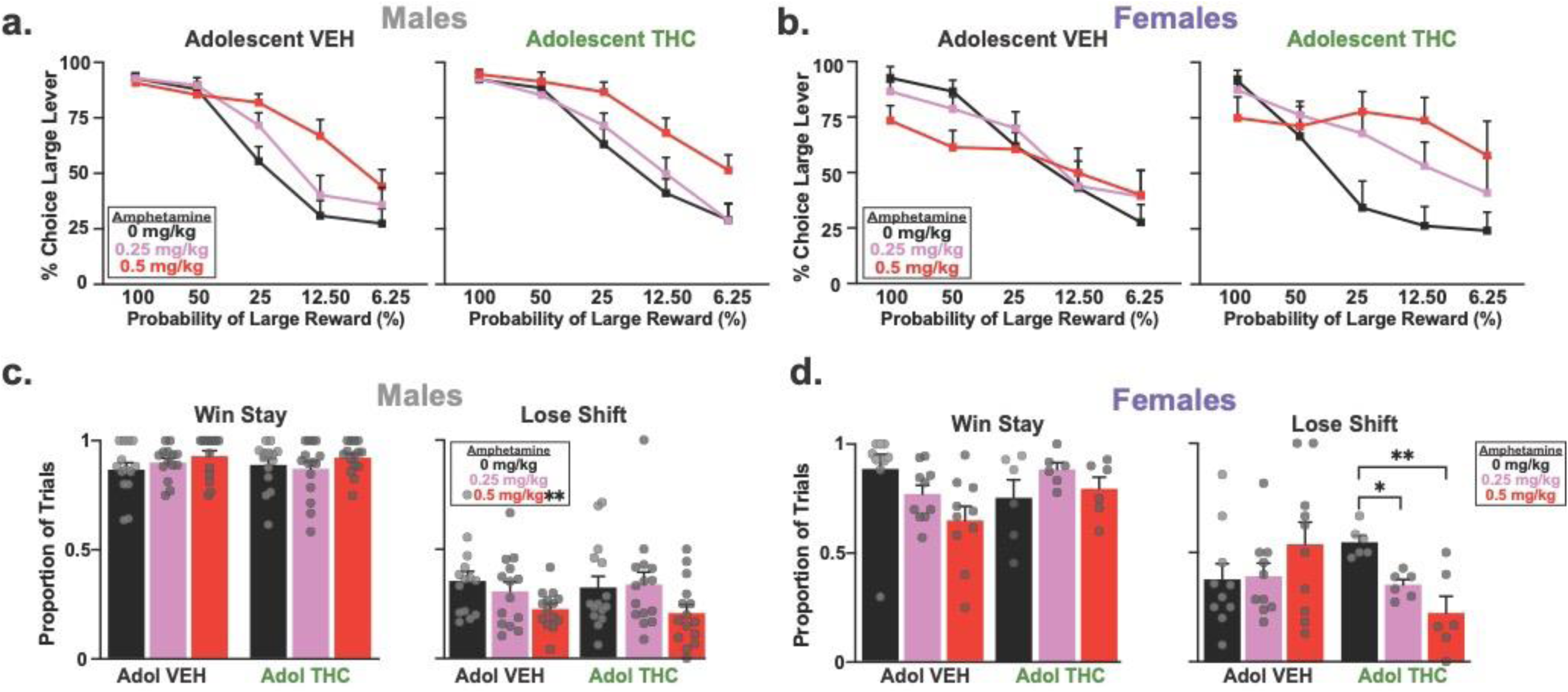
Amphetamine-induced ‘risky’ responding is potentiated after adolescent THC. **a)** Data from saline (black line/bar), low dose (0.25mg/kg; pink line/bar) and high dose (0.5mg/kg; red line/bar) amphetamine tests in each sex are shown. When performance patterns were further interrogated, we found that high dose of AMPH in Males increased inflexibility across both AdoTX groups, however in **b)** females, the high dose of AMPH in AdoVEH rats reduced risky responding at high probability blocks not seen in AdoTHC females. **c)** AMPH did not alter win-stay,but reduced lose-shift in AdoTX males. **d)** In AdoVEH females, AMPH decreased win-stay and increased lose-shift, while in AdoTHC AMPH had no effect on win-stay but decreased lose-shift. AMPH = amphetamine. AdoVEH (*n* = 14 males, 10 females), AdoTHC (*n* = 15 males, 6 females). Repeated measure three-way ANOVA; Sidak post hoc: *p\** < 0.05, *p\*\** < 0.01. Data represented as mean + SEM, individual animals shown as grey dots.

Subsequent analyses examined how amphetamine alters sensitivity to recent rewarded or non-rewarded choices by comparing win-stay and lose shift ratios. Analysis of the win-stay data yielded a significant amphetamine x AdoTX x Sex interaction (**Fig. 3c,d**, F_(2,82)_ = 4.34, *p* = 0.02). This was driven by the fact that in control rats, the 0.5 mg/kg dose resulted in lower win-stay values in females vs males (*p* < 0.01), although neither group showed significant changes in these values relative to saline (both Fs < 2.5, both *p*s > 0.10). Win-stay behavior was unaltered in AdoTHC rats (all *F*s < 2.4, all *p* > 0.10). In contrast, amphetamine had more pronounced effects on sensitivity to reward omissions, as indexed by changes in lose shift behavior. The analyses here revealed a significant amphetamine x AdoTX interaction (F_(2,38)_ = 4.84, *p* = 0.01) and a three-way interaction with the sex factor (F_(2,38)_ = 5.15, *p* = 0.01). In controls, amphetamine reduced lose shift behavior in males, but actually increased it in females (**Fig. 3c,d**, amphetamine x Sex: F_(2,44)_ = 3.86, *p* = 0.03, whereas in AdoTHC rats, these treatments uniformly reduced lose-shift behavior across sexes (amphetamine: F_(2,38)_ = 8.87, *p* < 0.001; amphetamine x Sex: F_(2,38)_ = 2.61, *p =* 0.08). From these data, we conclude that AdoTHC treatment makes rats more sensitive to the ability of amphetamine to increase risky choice and reduce sensitivity to losses, and this effect appears to be more prominent in females.

With respect to other performance measures, amphetamine increased choice latency and number of omissions across all probability blocks (amphetamine: Fs_(2,82)_ < 15.19, *ps* < 0.001, no amphetamine x Block interaction: Fs_(8,328)_ < 0.98, *ps* > 0.45). Analysis of the latency data revealed a significant amphetamine x AdoTX x Block (F_(8,328)_ = 2.27, *p* = 0.02) and amphetamine x AdoTX x Sex x Block interaction (F_(2,82)_ = 2.67, *p* = 0.01). Partitioning this latter interaction by AdoTX group revealed that in AdoVEH rats, females took longer to respond than males across the session (Sex x Block: F_(4,88)_ = 3.02, *p* = 0.02, Sex: F_(1,22)_ = 44.13, *p* < 0.001). In AdoTHC rat, females took longer to respond compared to males (Sex: F_(1,19)_ = 21.44, *p* < 0.001; no Sex X Block: F_(4,76)_ = 2.41, *p* = 0.06). Further analysis of the omission data revealed that overall females omitted more on trials compared to males (Sex x Block: F_(4,164)_ = 12.93, *p* < 0.001; Sex: Fs_(1,41)_ < 71.39, *p* < 0.001).

### Chemogenetic Dopamine Neuron Stimulation During Probabilistic Discounting

The above experiments showed that AdoTHC enhances visual cue learning in females but had few other effects on cognitive flexibility or probabilistic discounting under basal conditions. However, when treated with the monoamine-enhancing drug amphetamine, we found evidence for stronger enhancement of “risky” responding in AdoTHC rats, relative to AdoVEH controls. We therefore next asked whether this effect relates to changes in the functions of VTA dopamine neurons in particular, by using Gq-coupled DREADDs to acutely stimulate VTA dopamine neurons or VTA dopamine neuron projections to mPFC in rats with both AdoTX histories.

#### Initial Training

Rats in this experiment did not undergo strategy set shifting training prior to probabilistic discounting training, so we confirmed that AdoTX again did not affect initial acquisition of instrumental food seeking behavior during initial training (no main effect of AdoTX: Fs_(1, 46)_ < 0.78, *ps* > 0.38). We also confirmed that genotype (Geno: TH:Cre+ or WT littermate) did not alter instrumental training (Fs_(1,46)_ < 2.55, *ps* > 0.12; No AdoTX x Geno interaction: Fs_(1,46)_ < 0.63, *ps* > 0.43).

#### Probabilistic Discounting Training

Likewise, all included rats (TH:Cre and WT) successfully learned the discounting task (Main effect of Block: F_(4,184)_ = 129.26, *p* < 0.001, no main effects of, or interactions involving AdoTX or Geno: Fs_(1,46)_ < 0.88, *ps* > 0.35). Additionally, AdoTX nor genotype affected latency, omissions, or win-stay/lose-shift choice strategies (no main effects of, or interactions involving AdoTX or Geno: Fs_(1,46)_ < 3.34, *ps* > 0.07).

### Impact of Acutely Activating VTA Dopamine Neurons on Probabilistic Discounting

Despite the robust changes in locomotion and reward seeking that is seen following chemogenetic VTA dopamine neuron manipulations in TH:Cre rats (Boekhoudt et al. 2016; Boekhoudt et al. 2018; Halbout et al. 2019; Lawson et al. 2023; Mahler et al. 2019; Runegaard et al. 2018), we found few effects of VTA dopamine neuron stimulation on the probabilistic discounting task. First, we looked at VTA dopamine neuron stimulation in only TH:Cre rats. All rats displayed normal discounting profiles (TH:Cre+: main effect of Block: F_(4, 84)_ = 105.93, *p* < 0.001), and as in the prior experiment, no effect of AdoTX alone was seen on discounting (AdoTX x Block: F_(4, 84)_ = 0.91, *p* = 0.46). Moreover, when we tested the effects of systemic CNO to activate DA neurons, we did not observe any effects of this treatment in either control or AdoTX groups (**Fig. 4e**, no main effect of, or interactions involving AdoTX or CNO: Fs < 2.27, *ps* > 0.07). Additionally, we did not see any effects of CNO, AdoTX, nor interactions therein on choice latency, omissions, or win-stay/lose-shift choice strategy (Fs_(1,21)_ < 3.00, *ps* > 0.10; data not shown). These findings are inconsistent with our hypothesis that increased excitability of mesolimbic dopamine neurons would be sufficient to recapitulate the ability of amphetamine to increase risky/perseverative responding in either control or in AdoTHC-experienced rats.

**Fig. 4.**
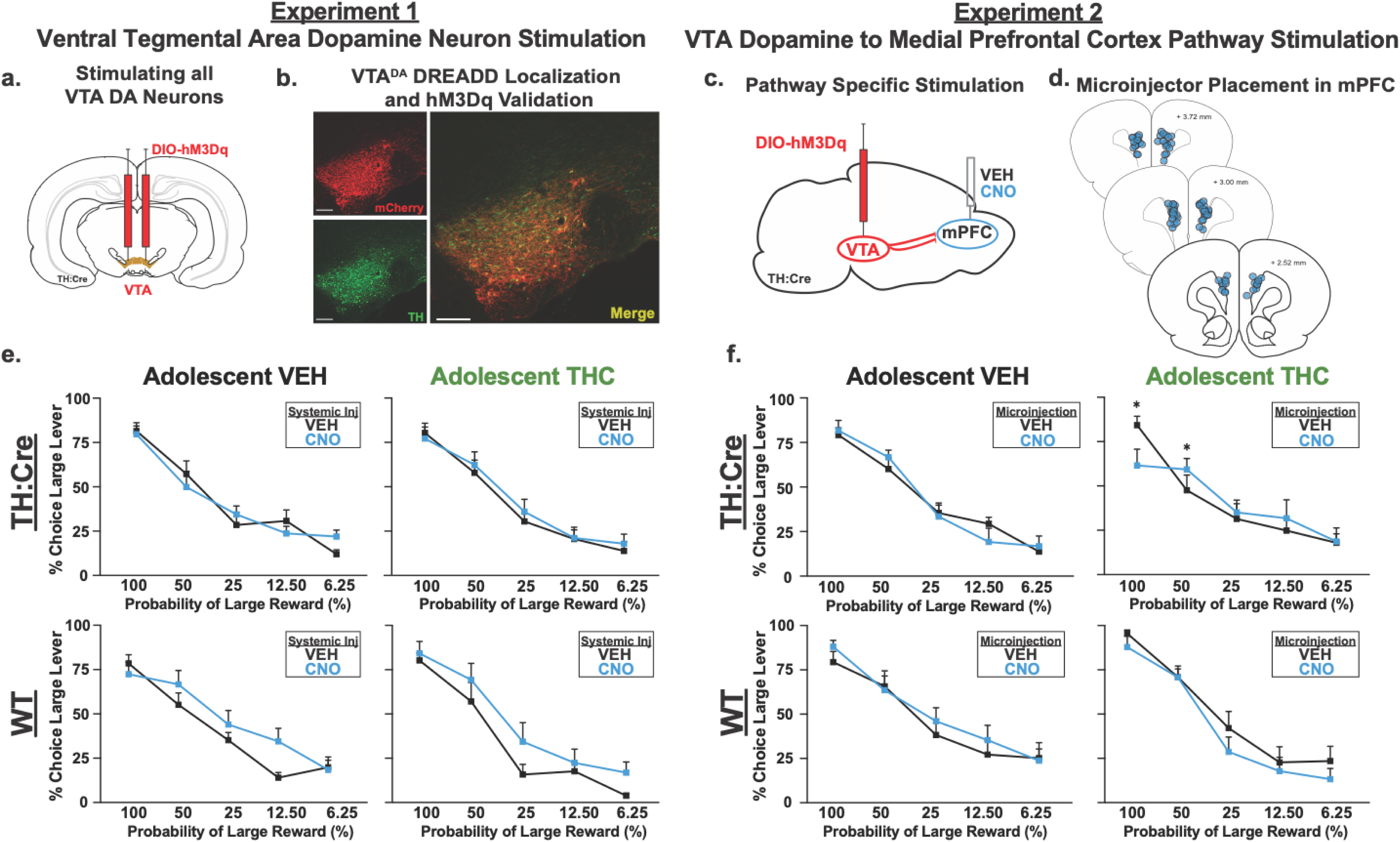
Stimulation of VTA dopamine neurons, or VTA dopamine projections to mPFC does not affect probabilistic discounting. **a)** Bilateral Cre-dependent hM3Dq DREADD AAV injections were made in VTA of TH:Cre rats, and of wildtype (WT) littermates. **b)** Example hM3Dq DREADD expression (red) is localized to tyrosine hydroxylase+ (TH; green) neurons within VTA (yellow=merge). Scale bar, 300 μm. **c)** For pathway-specific stimulation of VTA dopamine projections to mPFC, Cre-dependent hM3Dq DREADDs were injected into VTA as in Experiment 1, and cannulae targeting mPFC allowed CNO microinjection (1mM, 0.5 µl) upon DREADD-expressing dopamine neuron axons in this pathway. **d)** Cannula placements of each rat in pathway stimulation Experiment 2 is shown. **e)** Neither VTA dopamine neuron stimulation induced by systemic CNO in TH:Cre rats, nor **f)** stimulation of the VTA dopamine projection to mPFC induced by mPFC CNO microinjections in TH:Cre rats robustly altered probabilistic discounting in AdoTHC or AdoVEH rats. Likewise, lower panels of **e,f)** indicate that CNO did not have robust effects on WT rats without DREADDs. Data is represented as average % choice of the large reward lever in each probability block across the two CNO, and two VEH tests conducted in each rat, mean±SEM, *p\** < 0.05. TH:Cre+: VTA dopamine stimulation; AdoVEH (*n* = 8M, 3F) AdoTHC (*n* = 8M, 4F); VTA dopamine to mPFC: AdoVEH (*n* = 10M, 4F) AdoTHC (*n* = 9M, 5F). Wildtype: VTA dopamine stimulation; AdoVEH (*n* = 5M, 3F) AdoTHC (*n* = 5M); VTA dopamine to mPFC: AdoVEH (*n* = 9M, 4F) AdoTHC (*n* = 8M).

### Impact of Selectively Activating VTA Dopamine Projections to mPFC

We next asked whether selectively stimulating VTA dopamine neuron projections to mPFC would recapitulate the potentiation of amphetamine effects in AdoTHC rats. We did so by locally applying CNO upon DREADD-expressing axons of VTA dopamine neurons in mPFC. We have shown that this pathway-specific stimulation approach is capable of potentiating both axonal dopamine release and motivated reward seeking (Halbout et al. 2019; Mahler and Aston-Jones 2012; Mahler et al. 2019). We found few effects of this manipulation on probabilistic discounting. On the probabilistic discounting task, “risky” responding across the session varied across CNO and AdoTX (**Fig. 4**; AdoTX x CNO x Block interaction: F_(4,104)_ = 2.99, *p* = 0.02; no main effect of AdoTX: F_(1,26)_ = 0.15, *p* = 0.70; no main effect of CNO: F_(1,26)_ = 0.000 *p* = 0.10), in AdoVEH animals there were no measurable effects of or interactions with CNO (all Fs < 1.80, *ps >* 0.14). However, in AdoTHC rats CNO modestly reduced risky responding at the 100% probability block followed by an increase at the 50% probability block (CNO x Block: F_(4,52)_ = 2.99, *p* = 0.03). Rats performed similarly on choice latency, omissions, and win-stay/lose-shift regardless of AdoTX, CNO, or interactions therein with block (All Fs < 2.00, *p*s > 0.10).

### Minimal DREADD-independent Effects of CNO

Neither systemic nor intra-mPFC CNO had major behavioral effects in WT rats lacking DREADD expression (**Fig. 4e,f**). Systemic CNO in WT rats seemed to promote a more risk-prone phenotype, based on increased preference for the risky lever across all blocks (Main effect of CNO in WT rats: F_(1, 11)_ = 9.42, *p* = 0.01; no interactions of CNO, AdoTX, or Block: Fs_(4, 44)_ < 2.04, *ps* > 0.11), but did not affect response latency, omissions, or win-stay/lose-shift choice strategies (Fs_(1,11)_ < 2.96, *ps* > 0.11). Intra-mPFC CNO in WT rats altered risk responding such that “risky” choice was decreased in AdoTHC rats, whereas it was increased in AdoVEH animals (AdoTX x CNO: F_(1,16)_ = 5.40, *p* = 0.03). These effects were driven by changes in reward sensitivity, as CNO differentially affected win-stay behavior in the two groups (F_(1,160_ = 5.78, *p* = 0.03; no main effect of CNO F_(1,16)_ = 0.73, *p* = 0.41). Thus, CNO in AdoTHC WT rats were less likely to follow a risky win with another risky choice, while CNO had the opposite effects in in AdoVEH WT (data not shown). Additionally, we saw that AdoTHC animals had lower risky responding in the higher probability blocks, that then increased in the latter probability blocks (AdoTX x Block: F_(4,64)_ = 2.52, *p* = 0.05). We did not see any effect of AdoTX or CNO on lose-shift strategies (CNO x AdoTX: F_(1,16)_ = 0.67, *p* = 0.43; no main effect of CNO F_(1,16)_ = 1.71, *p* = 0.21) or response latency and omissions (Fs_(1,16)_ < 3.74, *ps* > 0.07; data not shown).

## Discussion

Here we show that administration of a well-characterized, human-relevant dose of THC (Ruiz et al. 2021a; Torrens et al. 2022; Torrens et al. 2020) during adolescence has subtle effects on behavioral tests of instrumental learning, without measurably altering performance on mPFC dopamine-dependent strategy set shifting or risk/reward decision making assessed with a probabilistic discounting task. However, when monoamine signaling was acutely enhanced with systemic amphetamine, an underlying effect of AdoTHC history was revealed. Relative to AdoVEH controls, amphetamine in AdoTHC rats were more sensitive to the ability of lower doses of amphetamine to increase perseverative responding for a large reward option when this choice was unlikely to result in reward. However, this potentiation of amphetamine effects in AdoTHC rats was not recapitulated by more specific chemogenetic stimulation of VTA dopamine neurons, or of their projections to mPFC in particular. This could suggest non-VTA dopaminergic mechanisms underlying potentiation of this cognitive effect of amphetamine in AdoTHC rats, suggesting potential impacts of THC on adolescent development of other systems upon which amphetamine acts. Alternatively, the enhanced effect of amphetamine reported here may be driven by alterations at the level of the dopamine terminal rather than changes in dopamine cell excitability, as dopamine receptor antagonism is able to block this amphetamine-induced risky responding (St. Onge and Floresco 2009). Further, we thoroughly characterize sex differences in decision-making across these behavioral tasks, some of which mediate the persistent impacts of AdoTHC on behavior. Results open new directions for investigating long-term impacts of AdoTHC on non-VTA dopaminergic modulation of cognition, and may inform associational studies of the long-term impacts of adolescent cannabis use in humans.

### Adolescent THC Exposure Enhances Initial Discrimination Learning

AdoTHC rats learned a visually cued instrumental response rule quicker than AdoVEH-treated controls, and this effect was most robust in females. This finding adds to the literature on potentially pro-learning/cognitive effects of AdoTHC (Hernandez et al. 2021; Stringfield and Torregrossa 2021b). This said, previous studies using other adolescent cannabinoid exposure models have not found analogous increases in initial rule discrimination learning (Freels et al. 2024; Gomes et al. 2015; Hernandez et al. 2021), though this might be due to the fact that few studies included female subjects, and the THC administration protocols employed were quite different.

We found no clear effects of AdoTHC across our set shifting performance metrics, including trials to criterion, number of errors made, or error types on the set shift day. These findings are consistent with others that found no major AdoTHC- induced changes in cognitive flexibility, as quantified in with different set shifting tasks, including the intra/extra dimensional “digging” task and an operant-based task similar to the one used here (Gomes et al. 2015; Hernandez et al. 2021; Poulia et al. 2021). However, in a study utilizing the same strategy set-shifting task as used here, females exposed to self-administered cannabis extract vapor during adolescence took longer to learn the new rule, and made more errors on the set-shift day, though this deficit was not seen after experimenter-administered cannabis vapor (Freels et al. 2024). It is presently unclear whether differences in patterns of results reflect differences in route of THC administration or dose, the specific timing of THC exposure during adolescence, or other experimental details. As such, this remains an important topic for future research intended to probe how adolescent cannabinoid exposure may alter executive functioning.

Though untested here, it is possible that AdoTHC altered other related cognitive processes such as transitioning between tasks, mental sets, or rule structures, which depend upon more lateral PFC subregions such as orbitofrontal cortex (OFC; Birrell and Brown 2000; Floresco et al. 2008; McAlonan and Brown 2003). For example, adolescent pubertal administration of the potent CBR agonist WIN 55,212 alters OFC-dependent reversal learning in and attentional set-shifting task (Gomes et al. 2015), though adolescent vaporized cannabis or cannabis smoke did not impact this same type of reversal learning (Freels et al. 2024; Hernandez et al. 2021). Again, discrepancies between studies may reflect differences in effects of cannabinoid drugs, doses, exposure timing, and washout period; further underscoring the need for a consistent, rationally-designed AdoTHC exposure model in the field—we argue that the model used here is the best-characterized to date in the field (Halbout et al. 2023; Lee et al. 2022; Lee et al. 2024; Lin et al. 2023; Ruiz et al. 2021a; Torrens et al. 2022; Torrens et al. 2020). This said, the possibility that OFC is even more sensitive to disruption by AdoTHC than mPFC should be directly tested in future studies.

### Adolescent THC Does Not Alter Basal Probabilistic Discounting

In the mPFC-dependent probabilistic discounting task (St. Onge and Floresco 2010), rats choose between a small reward that is always delivered when chosen (1 palatable banana pellet), and a larger reward that becomes increasingly unlikely to be delivered over the course of the ∼1h session (providing either 4 or 0 pellets). Efficient performance on this task demands evaluation of both risk and opportunity, and is dependent on intact functioning of both the mPFC and mesocortical and mesoaccumbens dopamine transmission (Jenni et al. 2017; St. Onge et al. 2012; St. Onge et al. 2010; Stopper et al. 2013). Impaired PFC activity results in deficits in adjusting choice in response to changes in reward probabilities, loss assessment, and a diminished ability to appropriately compare and favor larger rewards even when the probability of receiving them is higher (Bercovici et al. 2023; St. Onge and Floresco 2010). We found no major impacts of AdoTHC history on acquisition of, or stable performance on this task, as measured by choices of the ‘risky’ lever, choice strategies following rewarded vs unrewarded trials, latency to decide, and decisions to omit trials. One prior study (Jacobs-Brichford et al. 2019) found that administration of WIN 55,212 in both sexes during adolescence elevated preference for the ‘risky’ lever at lower reward probabilities, which we did not see in either sex as a result of our AdoTHC exposure model. Though several other differences exist between this study and the present one, we note that several prior reports have also shown more severe lasting effects of synthetic CBR agonists versus THC when administered in adolescents (Higuera-Matas et al. 2015; Renard et al. 2014; Stringfield and Torregrossa 2021a).

### Adolescent THC History Potentiates Amphetamine Effects on Probabilistic Discounting

Dopamine markedly influences mPFC-dependent cognition (Floresco and Magyar 2006; Goldman-Rakic 1995; Goto et al. 2007; Seamans and Yang 2004), including in the probabilistic discounting task employed here (Floresco and Whelan 2009; Islas-Preciado et al. 2020; Jenni et al. 2017; St. Onge et al. 2012; St. Onge et al. 2010; St. Onge and Floresco 2009). In the present study we replicated prior findings that pharmacologically challenging monoamine systems with amphetamine increased perseveration of responding for a large reward in both males and females (Islas-Preciado et al. 2020; St. Onge and Floresco 2009). This change was accompanied by a reduction in lose-shift behaviors following ‘risky’ losses, indicating that enhancing DA levels attenuates the impact that non-rewarded actions exert over subsequent choice. Interestingly, we saw that amphetamine led to an overall increase in choice latency, and omitted trials. This may have been driven in part by increased psychomotor activity induced by amphetamine that may have displaced animals from the levers, thus delaying choice.

Moreover, we found that this effect of amphetamine was more potent in AdoTHC, relative to AdoVEH rats, especially in females. At the highest dose of amphetamine, AdoTHC rats chose the ‘risky’ lever more when probabilities of that reward were low. These effects were also markedly sex-dependent. In males, amphetamine reduced overall lose-shift behavior. In females, the highest dose of amphetamine induced a disruptive reduction in win-stay behavior only in AdoVEH females, whereas it reduced lose-shift behaviors only in AdoTHC females, contributing to the heightened increased ‘risky’ responding observed in the AdoTHC groups. While we refer to this behavior as ‘risky’ responding (St. Onge et al. 2010), this behavior may also be interpreted as amphetamine causing AdoTHC animals to be less adaptive to changing contingencies, or more liable to perseverate on the initially prepotent large reward lever. Regardless, these results suggest that although probabilistic discounting under basal conditions is not altered by AdoTHC, underlying differences were nonetheless revealed upon acute activation of monoaminergic signaling. Amphetamine blocks and reverses the transporter for dopamine, norepinephrine, serotonin, and changes in one or more of these systems could be responsible for the potentiated response to amphetamine we see in this task in AdoTHC rats (especially females).

### Effects of Chemogenetic VTA Dopamine Neuron, and mPFC Dopamine Projection Stimulation on Probabilistic Discounting

In our next experiments, we sought to determine whether, as predicted, changes in VTA dopamine neuron signaling are responsible for the potentiated effect of amphetamine in AdoTHC rats. We therefore prepared transgenic TH:Cre rats with excitatory DREADDs, allowing stimulation of either all VTA dopamine neurons, or of dopamine neuron projections to mPFC in particular. Surprisingly, no major effects of chemogenetically stimulating all VTA dopamine neurons were found on probabilistic discounting. In this regard, it bears mentioning that pharmacological stimulation of dopamine D2 or D1 receptors in the nucleus accumbens (a main target of the VTA dopamine projection) either had no effect on risky choice or led to more optimal decision making, respectively-an effect distinct from that induced by amphetamine (Stopper et al. 2013). Furthermore, when we stimulated VTA dopamine neuron projections to mPFC, this did not markedly affect probabilistic discounting in AdoVEH rats, which is similar to a prior finding examining manipulations of VTA dopamine projections to PFC (Verharen et al. 2018). On the other hand, chemogenetic stimulation of mPFC dopamine terminals in AdoTHC rats attenuated risky responding during the 100% probability block, but then increased it in the subsequent 50% block. This effect loosely resembles that induced by intra-PFC D_2_ receptor stimulation on probabilistic discounting, which flattened the discounting curve (St. Onge et al. 2011). This is consistent with the possibility that AdoTHC exposure causes a slight enhancement in mesocortical dopamine D_2_ receptor signaling—a possibility that should be directly investigated. Moreover, no other differential effects of DREADD stimulation were seen in AdoTHC versus AdoVEH groups. The absence of significant effects in this experiment may also relate to the relatively small number of TH:Cre females, especially since amphetamine effects on the same behaviors seem to be driven by females. Additionally, the lack of behavioral effects of chemogenetic stimulation is unlikely to have resulted from inefficient DREADD stimulation or infection. Extent of viral expression, and selectivity of expression in TH+ VTA neurons was equivalent to our prior reports using these transgenic rats and viral vectors. In these reports we have shown this DIO-hM3Dq vector has >97% selectivity to TH+ neurons, and it is capable of driving dopamine neuron firing rates in vitro, increasing the number of them active in vivo, and increasing cFos expression in them (Mahler et al. 2019). Accordingly, we and others have found very robust behavioral effects of VTA dopamine neuron stimulation in a range of tasks, including cocaine intake and seeking, food reward intake, locomotion, and motivation (Boekhoudt et al. 2016; Boekhoudt et al. 2018; Halbout et al. 2019; Lawson et al. 2023; Mahler et al. 2019; Runegaard et al. 2018). We also previously confirmed that pathway-specific stimulation of VTA dopamine projections enhances dopamine release, and stimulation of mPFC projections is behaviorally efficacious for stimulating cue-induced cocaine seeking as well (Mahler et al. 2019). Therefore, it was particularly striking that we did not see any notable behavioral effects of dopamine neuron or dopamine projection stimulations on probability discounting, despite the fact that this task is sensitive to mPFC dopamine receptor manipulations. This may imply that alterations of VTA and non-VTA dopamine caused by amphetamine might simply be qualitatively distinct from manipulations of VTA dopamine neuron activity and dopamine release, as were conducted here chemogenetically (Mahler et al. 2019; Mahler et al. 2014). In this regard, performance of this probabilistic discounting task is associated with fluctuations in mesocortical and mesoaccumbens dopamine that relate to changes in the amount of rewards received and adjustments in risky choice biases (St. Onge et al, 2012). Amphetamine would cause robust increases in tonic dopamine levels, leading to a more static dopamine signal that impedes flexible adjustments in choice biases, whereas chemogenetic increases in dopamine neuron excitability would be a comparatively more subtle manipulation that could still permit variations in dopamine signaling over a session. Alternatively, amphetamine-induced changes in risk taking seen in AdoTHC rats may not solely depend upon VTA dopamine neurons themselves, but may instead depend upon other neural systems targeted by amphetamine, such as nigrostriatal dopamine projections to the dorsomedial striatum (Schumacher et al. 2021), other non-mPFC targets of VTA dopamine neurons, or other monoamine systems affected by amphetamine. Regardless, this intriguing finding refines our understanding of the neural substrates of probability-based decision making and underscores the need for further exploration to identify the specific mechanisms involved.

### Sex Differences

We conducted these studies in female as well as male rats, and found several consistent sex differences, both overall and in interaction with other experimental variables. Consistent with existing literature (Gargiulo et al. 2022; Islas-Preciado et al. 2020), we saw sex differences in response characteristics during operant set-shifting as well as probabilistic discounting, with females showing longer deliberation times (response latency), and more omitted trials than males. Strikingly, these specific sex differences consistently surfaced in the probabilistic discounting task across various experiments and cohorts, underscoring the robustness of the response latency and trial omission effects in females. Some of these sex differences may derive from females attaining satiety on the palatable reward quicker than males, but since both omissions and longer latencies at least partially emerged in females early in sessions during otherwise apparently vigorous reward seeking, this finding may also reflect different strategies taken by females relative to males (Chen et al. 2021; Orsini and Setlow 2017). For example, female rats have previously been shown to be more risk-averse than males on average (Orsini et al. 2016), and may exhibit prolonged response times and higher omission rates due to aversion to the risk of losing a reward, as demonstrated by a heightened sensitivity to loss (van den Bos et al. 2012). Alternatively, extended choice latencies and omissions may signify females’ tendency to take more time in learning about the probability distribution of the outcomes. This aligns with both rodent and human findings, indicating that females take longer to develop a preference for the more advantageous option when learning about probability distributions of reward versus punishment (van den Bos et al. 2013; van den Bos et al. 2012). Both possibilities warrant further exploration in future.

### Limitations

The present report has a number of limitations that should be considered. A single dose of THC (5mg/kg IP) was administered to adolescents daily from postnatal days 30 to 43, and persistent effects of THC are known to be dose-dependent (Amal et al. 2010; Freeman-Striegel et al. 2023; Irimia et al. 2015), and dependent upon the specific developmental stage at which it is experienced (Cha et al. 2006; Gorey et al. 2019; Mokrysz et al. 2016; Murray et al. 2022; Schramm-Sapyta et al. 2007; Torrens et al. 2022; Torrens et al. 2020). The low-dose THC regimen used here may capture the effects of frequent, low-potency cannabis use (Cooper and Haney 2009; Huestis et al. 1992) rather than high-intensity or chronic cannabis exposure, use of high-potency cannabis products, or escalating use over time. Further work should examine how THC dose, dosing pattern, and means of administration (i.e. oral vs intraperitoneal vs inhalation) impact long-term function of dopamine-dependent cognition. Also relevant to dose, we and others often observe more potent behavioral effects of THC in female, relative to male adolescent rats, which is likely attributable in part to differences in THC metabolism, in particular with regard to a higher blood and brain levels in female adolescent and adult rats and mice of the THC metabolite 11-OH-THC, a CB1 agonist (Baglot et al. 2021; Craft et al. 2019; Ruiz et al. 2021a; Ruiz et al. 2021b; Torrens et al. 2022; Torrens et al. 2020; Tseng et al. 2004; Wiley and Burston 2014), and this sex difference is also seen in humans after cannabis use (Arkell et al. 2022; Matheson et al. 2020; Sholler et al. 2021). This metabolic sex difference may at least in part account for the more robust persistent effects of adolescent THC in females than males seen here, and in numerous prior reports (Cha et al. 2007; Le et al. 2022; Rubino and Parolaro 2011; Rubino et al. 2009; Ruiz et al. 2021a; Ruiz et al. 2021b; Tseng et al. 2004). We found that in females, AdoTHC had more pronounced effects than in males, including enhancement of visual cue learning, and stronger potentiation of amphetamine’s effects on probability discounting behavior—but no effect on other behaviors like set shifting and baseline probability discounting. These results support the increasingly apparent possibility that AdoTHC’s effects are both highly sex- and task-dependent. Further studies investigating the reasons why AdoTHC seems to influence some behaviors and not others are needed, and may provide important new evidence relevant to cognitive and anatomical development. Another limitation of this report is the lack of data on complementary cognitive tasks, which are dependent upon other cortical regions like OFC that are notably sensitive to adolescent cannabinoid exposure (Egerton et al. 2005; Gomes et al. 2015; Klugmann et al. 2011), and which are mediated by ECB-dependent processes that may be particularly sensitive to AdoTHC’s long-lasting impacts.

### Summary

In sum, this paper provides novel information about the impact of AdoTHC on the developing brain of males, and especially of females. Results point to long-lasting changes that influence cue learning, and the ability of acute amphetamine to alter risky decision making. A key finding is the pervasive nature of sex-differences—both in behavioral strategies employed by rats in these tasks, and in the severity of long-lasting consequences of AdoTHC.

