## Supplemental Figure for "Adolescent THC impacts on mPFC dopamine-mediated cognitive processes in male and female rats"

**Fig. S1. Stimulation of VTA dopamine neurons, or VTA dopamine projections to mPFC does not affect probabilistic discounting in either sex** a) VTA dopamine neuron stimulation induced by systemic CNO in TH:Cre AdoTX female and male rats b) stimulation of the VTA dopamine projection to mPFC induced by mPFC CNO microinjections in TH:Cre AdoTX female and male rats. TH:Cre+: VTA dopamine stimulation; AdoVEH (*n* = 8M, 3F) AdoTHC (*n* = 8M, 4F); VTA dopamine to mPFC: AdoVEH (*n* = 10M, 4F) AdoTHC (*n* = 9M, 5F).

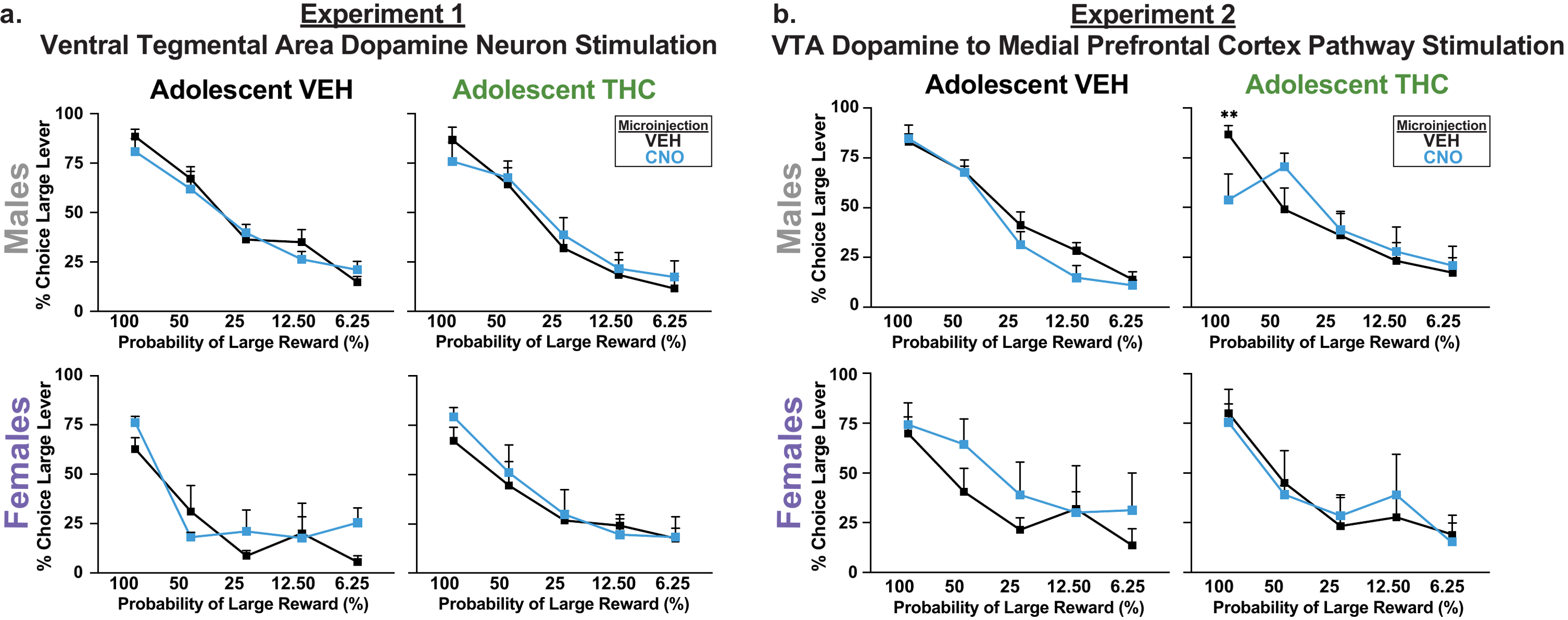
